# MiR-221/222-enriched ADSC-exosome mitigates PM exposure-exacerbated cardiac ischemia/reperfusion injury through the modulation of the BNIP3/LC3B/PUMA pathway

**DOI:** 10.1101/2023.11.10.566671

**Authors:** Tzu-Lin Lee, Wen-Chi Shen, Ya-Chun Chen, Tsai-Chun Lai, Shu-Rung Lin, Shu-Wha Lin, I-Shing Yu, Yen-Hsiu Yeh, Tsai-Kun Li, I-Ta Lee, Chiang-Wen Lee, Yuh-Lien Chen

## Abstract

**Background:** Epidemiology has demonstrated a strong relationship between fine particulate matter (PM) exposure and cardiovascular disease. Whether PM aggravates myocardial ischemia/reperfusion (I/R) injury and its related mechanisms remain unclear. Our previous study showed that adipose stem cell-derived exosomes (ADSC-Exo) contain a large amount of miR-221/222. This study investigated the effects of PM exposure on I/R-induced cardiac injury through mitophagy and apoptosis, as well as the potential role of miR-221/222 in ADSC-Exo.

**Methods:** Wild-type, miR-221/222 knockout (miR-221/222 KO), and miR-221/222 overexpressed transgenic (miR-221/222 TG) mice were intratracheally injected with 100 μg/kg PM for 24 h before I/R treatment. Ischemia was induced by temporarily occluding the left anterior descending (LAD) coronary artery with sutures for 30 min, followed by 3 h of reperfusion. In an *in vitro* model, H9c2 cells were exposed to 50 μg/mL PM for 6 h and subjected to hypoxia (1% O_2_) at 37°C for 6 h, followed by 12 h reoxygenation.

**Results:** PM aggravates I/R (H/R)-induced cardiac injury by increasing ROS levels and causing mitochondrial dysfunction, leading to an increase in mitochondrial fission-related proteins like Drp1 and Mff, mitophagy-related proteins such as BNIP3 and LC3B, as well as apoptosis-related proteins like PUMA and p-p53 *in vivo* and *in vitro* studies. In comparison, transfection of ADSC-Exo and miR-221/222 mimics significantly reduced PM+I/R (H/R)-induced cardiac injury. Importantly, ADSC-Exo contains miR-221/222, which directly targets BNIP3, LC3B, and PUMA, decreasing their expression and ultimately reducing cardiomyocyte mitophagy and apoptosis.

**Conclusions:** The study showed that PM aggravates I/R or H/R-induced cardiac injury, and ADSC-Exo treatment significantly reduced this by regulating mitophagy and apoptosis through miR-221/222/BNIP3/LC3B/PUMA.

## Introduction

Growing epidemiological evidence suggests that short- and long-term exposure to particulate matter (PM), especially fine particulate matter (PM2.5), can lead to the induction, progression, and worsening of cardiovascular disease (CVD). ^1^ Inhalation of PM2.5 causes oxidative stress and inflammatory responses not limited to the lungs. However, it can also reach the systemic circulation and related organs/tissues (e.g., heart, vasculature, adipocytes). Evidence from animal models suggests that atherosclerosis, hypertension, thrombosis, and heart failure associated with cardiovascular disease may be exacerbated by PM2.5. ^2,3^ Coronary artery disease is the most common form of CVD. The medical strategy is timely reperfusion, such as percutaneous coronary intervention (PCI) and coronary artery bypass grafting (CABG). ^4^ However, while reperfusion promotes blood flow recovery in the border zone of myocardial infarction and rescues myocardial cells, it also aggravates cardiac ischemia/reperfusion (I/R) myocardial injury. It has increased mortality over the past few decades. Efforts have been made to elucidate the precise cellular and molecular mechanisms of cardiac I/R injury, such as the interaction between autophagy and apoptosis. ^5^ However, PM-related I/R cardiac injury and its associated mechanisms, especially mitochondrial function, remain poorly understood.

Several mechanisms, including reactive oxygen species (ROS) overproduction, apoptosis, and imbalance of mitochondrial dynamics, have been proposed to contribute to cardiac I/R injury. ^6^ The previous study reported that increased mitochondrial fission and decreased mitochondrial fusion were observed during cardiac I/R injury, which resulted in cardiac damage. ^7^ Mitochondrial fission is regulated by dynamin-related protein 1 (Drp1) and mitochondrial fission factor (Mff). Furthermore, mitochondria are critical for maintaining normal cellular functions through mitophagy. ^8^ The role of mitophagy remains controversial. Mitophagy has both beneficial and adverse effects on CVD. ^9^ Autophagy is thought to be cardioprotective during ischemia but detrimental during reperfusion. ^10,11^ BNIP3 mediates mitophagy through its cytoplasmic LIR granules, interacting with LC3 family proteins. ^12^ In addition, myocardial I/R injury induces JNK activation, increasing Mff and BNIP3 activity, promoting excessive mitochondrial fission, and mitophagy, as well as leading to cell death. ^13^ BNIP3 is activated upon ischemic stimulation and promotes cardiomyocyte death. ^14^ On the other hand, PM2.5 induced oxidative stress, leading to DNA damage, mitochondrial swelling, autophagy, and apoptosis in HaCaT cells and mouse skin tissue. ^15^ Furthermore, PM increased ROS levels and triggered cardiomyocyte apoptosis in AC16 cardiomyocytes. ^16,17^ Short-term exposure to PM triggered mitophagy, leading to mitochondrial dysfunction and partially mediating vascular fibrosis. ^18^ There are currently no reports elucidating the correlation between myocardial damage, disruption of the fission-fusion process, and mitophagy caused by the combined treatment of PM and I/R or H/R.

Adipose-derived stem cells (ADSCs) secrete paracrine products, including cytokines, growth factors, RNA, and microRNA, which can be delivered to target organs for repair. ^19^ As an essential transporter of paracrine factors, exosomes play an important role in angiogenesis, immune regulation, and tissue regeneration. ^20^ Our previous studies have shown that exosomes from ADSCs can protect flaps and the heart from ischemia/reperfusion injury. ^21,22^ There is much evidence that miRNAs are involved in various biological regulations, including myocardial mitochondrial ROS, inflammation, and apoptosis. ^23,24^ Some evidence suggests that miR-221/222 is a key regulator of cardiovascular function and tissue metabolism. ^25^ In addition, miR-221/222 can promote migration and invasion and inhibit autophagy and apoptosis. ^26^ On the other hand, miR-222 was identified to be associated with particulate matter exposure alone. ^27^ In this study, we used miRTarBase to predict the specific binding sites of miR-221/222 in the 3’ untranslated regions of BNIP3 and LC3B. Therefore, miR-221/222 may regulate mitophagy by regulating BNIP3 and LC3B genes. Currently, there is limited knowledge regarding the involvement of miR-221/222 in the induction of cardiomyocyte apoptosis and mitophagy when PM and I/R (or H/R) are combined treatments. This study demonstrates that PM aggravates cardiac injury caused by I/R or H/R. MiR-221/222 enriched in ADSC-Exo affects ROS production, mitochondrial dynamics, and mitophagy, and reduces cardiac injury through the miR-221/222/BNIP3/LC3B/PUMA pathway.

## Materials and Methods

Detailed procedures are described in the Supplemental Material.

## Result

### PM exposure aggravates I/R-induced cardiac injury, including ROS production and mitochondrial dynamics *in vivo*

First, whether PM affects cardiac function in mice under I/R conditions was assessed by echocardiography. Representative echocardiographic images were obtained from control, PM, I/R, and PM+I/R treated mice. Compared with controls, mice treated with PM or I/R alone had significantly reduced ejection fraction (EF) and fractional shortening (FS) percentages, indicating impaired systolic contractility. More importantly, the PM+I/R group exhibited even lower EF and FS compared to those treated with PM or I/R alone (Figure 1A). In addition, the plasma levels of lactate dehydrogenase (LDH) and troponin I, indicators of cardiac injury, were significantly elevated in the PM+I/R group (Figure 1B). Analysis of the ischemic area using TTC staining showed that the PM+I/R group significantly increased the white infarct area compared to those treated with PM or I/R alone (Figure 1C). Further, we investigated the underlying mechanisms affecting cardiac function impairment caused by PM exposure in I/R-treated mice. The proportion of apoptotic cells was significantly increased in PM+I/R compared with PM or I/R alone using the TUNEL assay (Figure 1D). We also found that the expression of PUMA, p-p53, and cleaved caspase 3 was significantly increased. In contrast, the expression of BCL2 (an anti-apoptotic protein) was decreased after PM+I/R treatment (Figure 1E). I/R-induced cardiac injury is often caused by ROS production. ^28^ Excessive ROS production resulted in mitochondrial dysfunction. Dihydroethidium (DHE) and MitoSOX Red are selectively used to measure cell and mitochondrial ROS, respectively. ^29^ The results showed that PMs with I/R operation exhibited substantial ROS accumulation in the cardiac section (Figures 1F, 1G). Furthermore, TEM analysis of heart tissue revealed mitochondrial ultrastructural changes in the hearts of PM+I/R-treated mice. In the control group, there were many normal mitochondria and myofilaments and few autophagosomes in the cytoplasm of cardiomyocytes. In contrast, the PM+I/R group showed many autophagosomes and mitochondrial fission (Figure 1H). Further, to gain a more comprehensive understanding of mitochondrial dysfunction in PM+I/R hearts. We assessed the mitochondrial fission-related proteins Drp1 and Mff and the mitophagy-related proteins Beclin-1, BNIP3, and LC3B levels by Western blot. The protein expression of Drp1, Mff, Beclin-1, BNIP3, and LC3B were significantly increased in the PM+I/R compared with the PM and I/R alone (Figure 1I). These results elucidate that exposure to PM exacerbated I/R-induced cardiac diseases, increased ROS production, and accelerated apoptosis and mitophagy.

**Figure 1:**
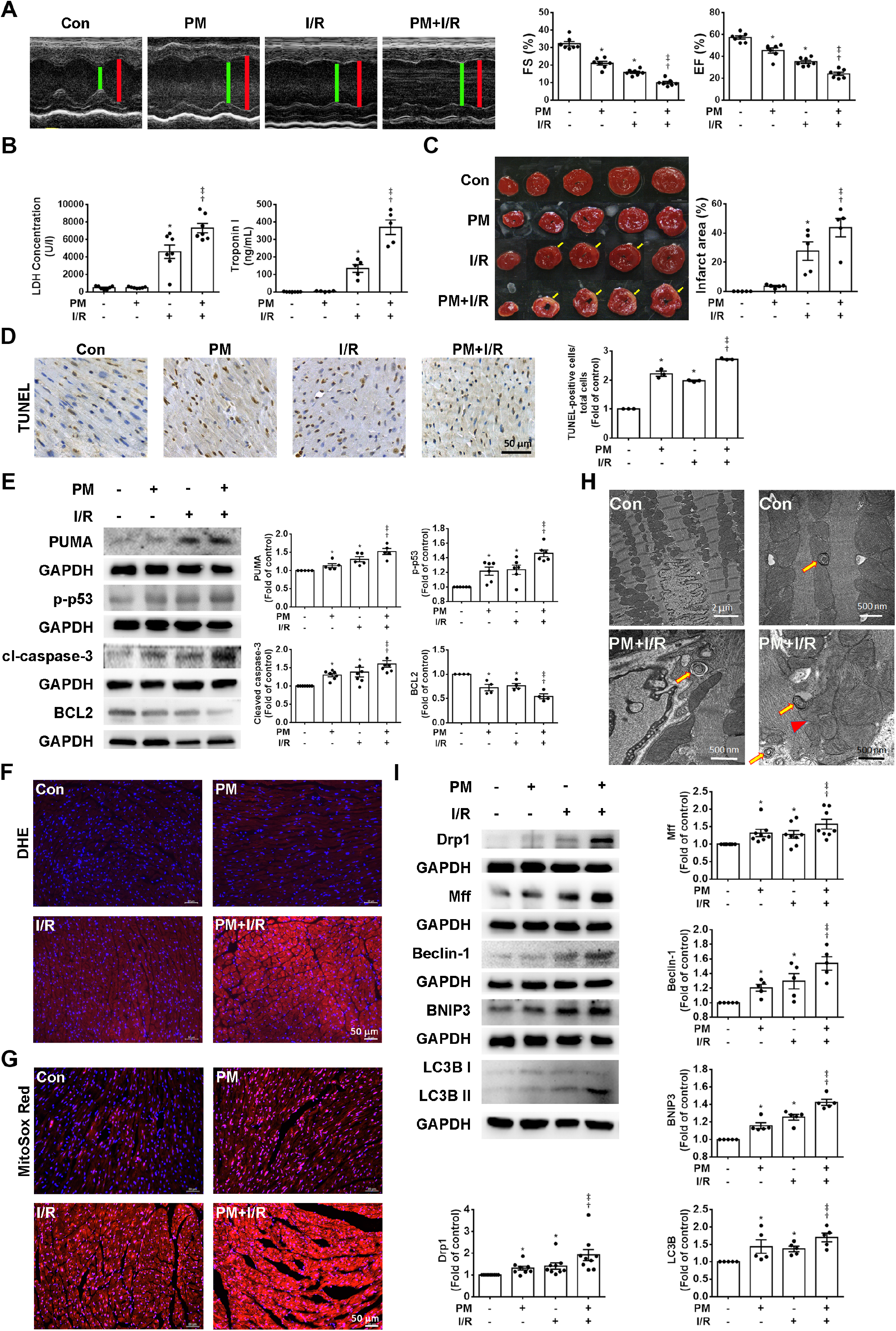
PM significantly impairs cardiac function and increases apoptosis in I/R-treated hearts of C57BL/6J mice. C57BL/6J mice were intratracheally injected with PM for 24 h, followed by 30 min of ischemia and 3 h of reperfusion. (A) The images of left ventricular end-systolic diameter (green line) and left ventricular end-diastolic diameter (red line) were performed by echocardiography. FS and EF percentages were measured in control, PM, I/R, and PM+I/R mice. (B) Cardiac injury was assessed by measuring plasma LDH and troponin I levels. (C) TTC staining was used to detect the ischemic area (scale bar=10 mm). The yellow arrows indicated the ischemia area. (D) Apoptotic cells were assessed by TUNEL assay (brown). Nuclei were counterstained by hematoxylin staining (blue). Scale bar=50 μm. (E) Western blot detects the expression of apoptosis-related proteins (PUMA, p-p53, cl-caspase-3, and BCL2). (F, G) Intracellular and mitochondrial ROS were measured using DHE and MitoSox Red respectively, and nuclei were stained blue with DAPI (scale bar=50 μm). (H) Ultrastructural morphology observed under the TEM. Autophagosome and mitochondrial fission are shown by arrowheads and arrows, respectively. (I) Western blot examined the expression of mitochondrial fission-related proteins (Drp1, Mff) and mitophagy-related proteins (Beclin-1, BNIP3, and LC3B). Data are expressed as mean ± SEM (n=3–8). *P < 0.05 compared with C, †P < 0.05 compared with PM, ǂP < 0.05 compared with I/R.

### PM exacerbates cardiomyocyte apoptosis through ROS generation under H/R conditions

To investigate the detailed mechanism by which PM exacerbates I/R-induced cardiac injury, we established a cell model with PM concentration and incubation time of 6 h, followed by H/R condition (6 h/12 h). In the H/R model, cell viability decreased after 6 h of pretreatment of H9c2 cells with 10 or 50 µg/mL PM (Figure 2A). Cells were treated with PM (50 µg/mL) and subjected to H/R (6 h/12 h) to evaluate the apoptosis of H9c2 cells by TUNEL assay and Annexin V-FITC/PI assay. TUNEL analysis showed that apoptosis was significantly increased in the PM+H/R group compared with the PM or H/R group alone (Figure 2B). The Annexin V-FITC/PI assay showed that PM+H/R treatment accelerated apoptosis (Figure 2C). Furthermore, the PM+H/R group had significantly increased PUMA, p-p53, and cleaved caspase 3 expression. It significantly decreased BCL2 expression compared with the group receiving PM or H/R alone (Figure 2D). To study the effect of PM+H/R-treated H9c2 on mitochondrial ROS. MitoSox Red and TO-PRO-3 double staining were used to examine the production of mitochondrial ROS. The results showed higher in living cells (MitoSox Red positive and TO-PRO-3 negative) under PM+H/R conditions (Figures 2E, 2F). Furthermore, as shown by fluorescence microscopy and flow cytometry, PM+H/R treatment significantly increased cytoplasmic H_2_O_2_ in living cells (PI negative and DCFH-DA positive) (Figures 2G, 2H). Likewise, the DHE overlay showed that PM+H/R increased the production of cytoplasmic superoxide anions (DHE-positive and DiOC_6_(3)-positive) as measured by fluorescence microscopy and flow cytometry (Figures 2I, 2J). The addition of MitoTEMPO, a mitochondrial antioxidant, inhibited PM+H/R-induced mitochondrial ROS expression by fluorescence microscopy and flow cytometry (Figure 2K, 2L). Western blot and TUNEL assay showed that MitoTEMPO treatment reduced PUMA and p-p53 expression while increasing BCL2 expression and reducing apoptotic cells (Figures 2M, 2N). These results indicated that PM+H/R treatment increased ROS production *in vitro*, leading to apoptosis and cardiac damage.

**Figure 2:**
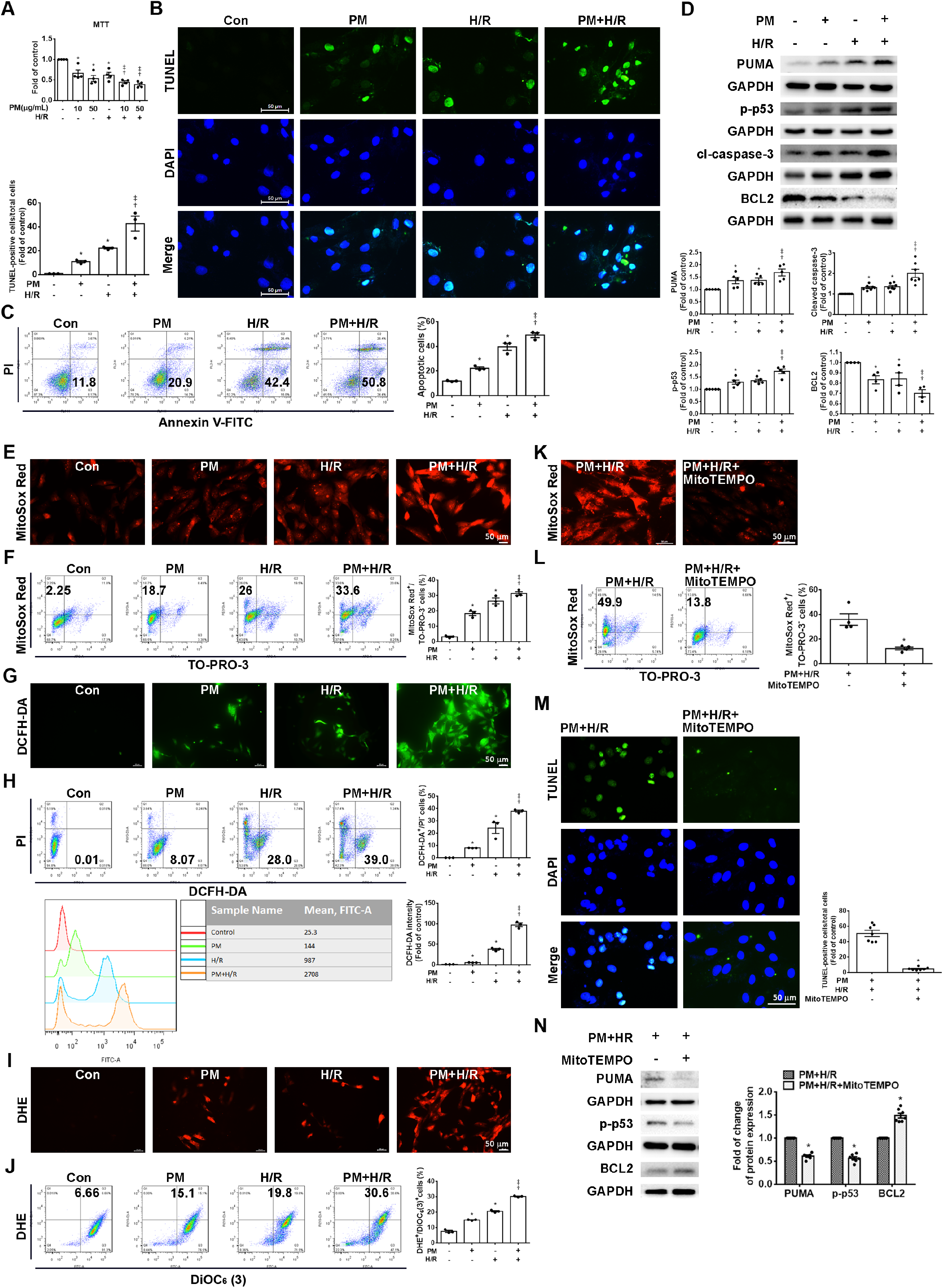
PM significantly exacerbated H/R-induced cardiomyocyte damage. H9c2 cells were pretreated with or without different concentrations of PM (10 or 50 μg/mL) for 6 h, followed by treatment with or without hypoxia for 6 h and reoxygenation for 12 h. (A) Cell viability was assessed by MTT assay. PM (50 μg/mL) was used in the following experiments. (B) TUNEL assay was used to assess apoptosis. Nuclei were stained by DAPI staining (blue color) and TUNEL-positive cells (green color). (C) Flow cytometry was used to measure apoptotic cells with PI/FITC-labeled annexin V staining. (D) The levels of apoptosis-related proteins were measured using Western blot. (E) Fluorescence microscopy was used to detect ROS production by MitoSox Red (1 μM) staining. (F) MitoSox Red (1 μM)/TO-PRO-3 (100 nM) staining was used to identify viable cells producing mitochondrial ROS (MitoSox Red-positive/TO-PRO-3-negative) by flow cytometry. (G) By fluorescence microscopy, DCFH-DA (10 μM) was used to detect cytoplasmic H_2_O_2_. (H) DCFH-DA/PI staining was used to examine viable cells producing cytoplasmic H_2_O_2_ (DCFH-DA-positive/PI-negative) by flow cytometry. (I) DHE (5 μM) was used to detect cellular superoxide anions by fluorescence microscopy. (J) DHE/DiOC_6_(3) (90 nM) staining was used to examine viable cells producing cellular superoxide anions (DHE-positive/DiOC_6_(3)-positive) by flow cytometry. (K-N) H9c2 cells were pretreated with 10 nM MitoTEMPO, an inhibitor of ROS production, for 1 h before exposure to PM and H/R. MitoSox Red is used to detect mitochondrial ROS production through fluorescence microscopy and flow cytometry. The TUNEL assay and Western blot detected the degree of cell apoptosis. Scale bar=50 μm. Data are expressed as mean ± SEM (n=3–7). *P < 0.05 compared with C; †P < 0.05 compared with PM; ǂP < 0.05 compared with H/R.

### PM exacerbates apoptosis in H/R-treated cardiomyocytes by worsening mitochondrial function and increasing mitophagy

Previous studies have demonstrated that impaired mitochondrial function, or excessive mitochondrial division can lead to cell death and cardiac injury. ^30,31^ ATP concentration, oxygen consumption rate (OCR), and mitochondrial membrane potential (ΔΨm) are critical parameters of mitochondrial function. ATP concentration was significantly reduced in PM or H/R treatment compared to the control, and the reduction was more significant in PM+H/R combined treatment compared to PM or H/R treatment alone (Figure 3A). OCR from the Seahorse metabolic analyzer was used to assess mitochondrial function. Compared with the control, PM or H/R significantly decreased OCR (maximal) and increased ATP, indicating that PM and H/R inhibited oxygen consumption and promoted extracellular acidification (Figure 3B). We determined the effect of PM+H/R on the changes in ΔΨm by JC-1 assay. PM+H/R caused a marked difference in the transition of fluorescence emission from red to green, which indicated a dissipation of ΔΨm (Figure 3C). Furthermore, flow cytometric analysis of JC-1 showed that PM+H/R treatment increased the number of low ΔΨm cells (JC-1 green positive and red negative) (Figure 3D). MitoTracker was used to examine the effect of PM+H/R on mitochondrial fission. The length of mitochondria in PM+H/R treated cells was significantly shorter than in other treated cells (Figure 3E). To further investigate how PM+H/R regulates mitochondrial fission, the levels of Drp1 and Mff were examined using Western blot. Our results showed that the expression of Drp1 and Mff was remarkably increased in the PM or H/R group compared with the control group. Furthermore, PM+H/R treatment significantly enhanced the expression of Drp1 and Mff (Figure 3F). Additionally, the effect of PM on mitophagy in H9c2 cells under H/R treatment was verified by acridine orange (AO) staining. ^32^ PM+H/R-treated cells showed an increase in the presence of autolysosomes (Figure 3G). To investigate how PM+H/R affects mitophagy, we analyzed the expression of Beclin-1, BNIP3, and LC3B by Western blot. PM+H/R increased the expression of mitophagy-related proteins (Figure 3H). We then investigated whether PM+H/R affects the translocation of Drp-1, Mff, Beclin-1, BNIP3, and LC3B from the cytoplasm to mitochondria in cardiomyocytes. PM+H/R significantly increased the expression of these proteins in mitochondria compared with PM- or H/R-treated alone cells (Figure 3I). Furthermore, fluorescent staining showed that PM increased the localization of BNIP3 and LC3B in the mitochondria of H/R-treated cells (Figure 3J). In addition, normal mitochondria were present in the cardiomyocytes’ cytoplasm of the control group, while several autophagosomes and PM particles were observed in the PM+H/R group (Figure 3K). Furthermore, Mdivi-1, a mitochondrial fission inhibitor, reduced PM+H/R-induced mitochondrial dysfunction, as shown by the JC-1 assay (Figures 3L, 3M). PM+H/R combined treatment increased autolysosomes, while Mdivi-1 treatment inhibited red fluorescence by AO staining (Figure 3N). Bafilomycin A1 is commonly used as an inhibitor of autophagosome-lysosome fusion to measure autophagic flux activity ^33^. Mdivi-1 and Bafilomycin A1 significantly reduced PM+H/R-induced apoptosis, as shown by the TUNEL assay (Figure 3O). PM+H/R exposure increased the Drp1, Mff, LC3B, BNIP3, PUMA and p-p53, but decreased BCL2. However, the Mdivi-1 treatment reversed the expression of these proteins (Figure 3P). Similarly, Bafilomycin A1 treatment reversed the PM+H/R-affected expression of mitophagy- and apoptosis-related proteins (Figure 3Q). These findings indicate that PM+H/R treatment affects mitochondrial function, mitophagy, and mitochondrial fission, suggesting a potential role for these mechanisms in the cellular response to PM+H/R-induced cardiac injury.

**Figure 3:**
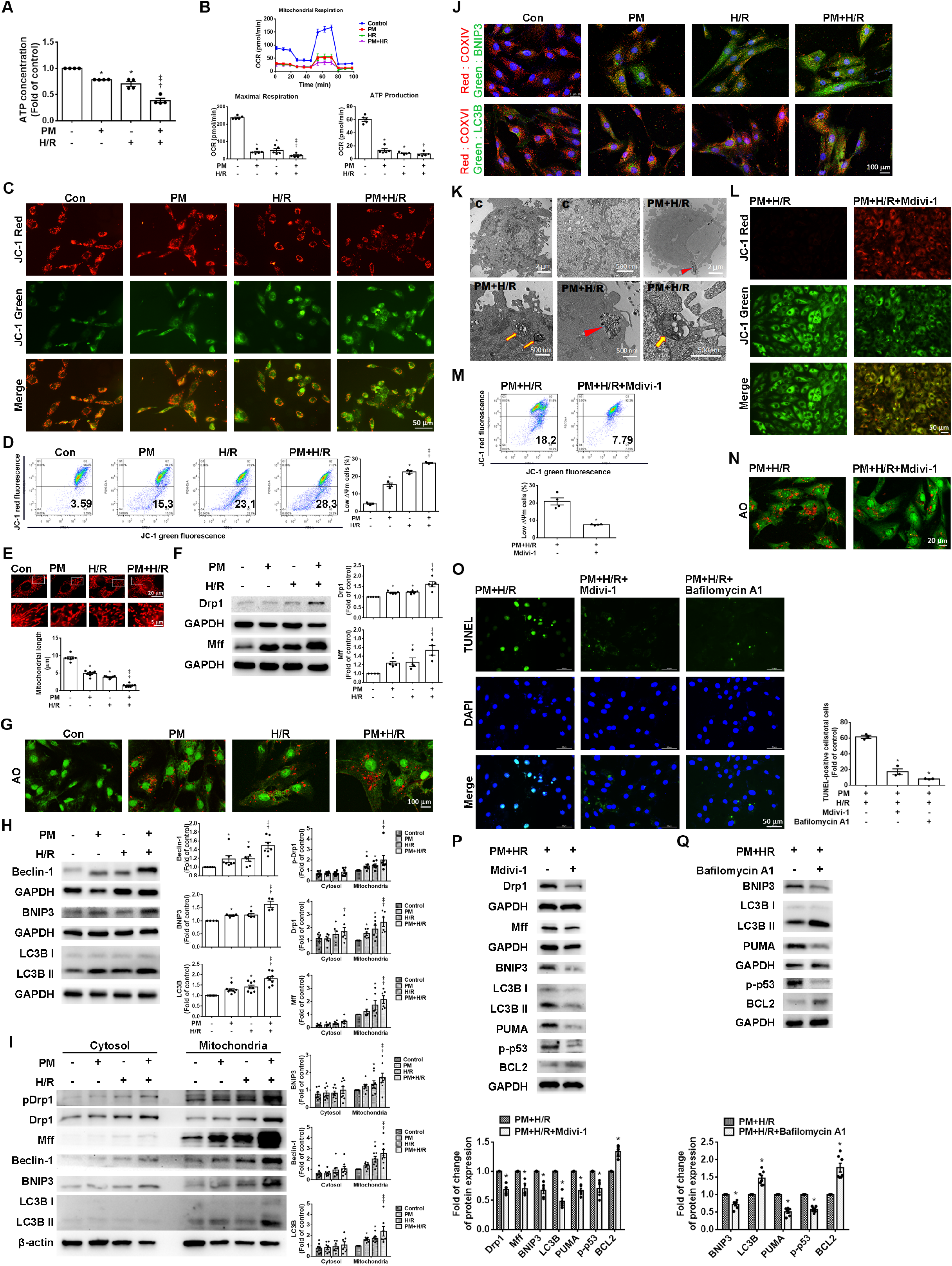
PM exacerbated the effects on mitochondrial dysfunction of H/R treated cardiomyocytes. Cardiomyocytes were pretreated with or without 50 μg/mL PM and then exposed to hypoxia for 6 h, followed by reoxygenation for 12 h. (A) Quantitative the ATP levels were detected. (B) OCR was determined using a Seahorse metabolic analyzer. Cells were stimulated sequentially with 1 μM oligomycin, 0.5 μM FCCP, and a mixture of 1 μM antimycin A and Rotenone to assess the mitochondrial stress tolerance. Quantitative results of ATP levels and maximal OCR were analyzed and shown. (C, D). The ΔΨm was determined by JC-1 staining under the fluorescent microscope and flow cytometry. High ΔΨm and low ΔΨm are shown in red and green, respectively. Scale bar=50 μm. (E) Mitochondrial length was measured with MitoTracker staining. The photo below is an enlargement of the rectangle above. Scale bar=20 or 5 μm, as indicated in the panel. (F) The levels of Drp1 and Mff were detected by Western blot. (G) Autolysosomes were observed with AO staining under a fluorescence microscope. Scale bar=100 μm. (H) Beclin-1, BNIP3, and LC3B expression levels were examined by Western blot. (I) The levels of pDRP-1, DRP-1, Mff, Beclin-1, BNIP3, and LC3B in the cytoplasmic and mitochondrial fractions were examined by Western blot. (J) The co-localization of BNIP3 (green) or LC3B (green) and COXIV (mitochondria, red) was observed by immunofluorescent microscopy. Scale bar=100 μm. (K) The ultrastructure of PM and mitophagosome was observed using TEM, as indicated in the panel. Bar=2 μm or 500 nm. (L, M) The effect of Mdivi-1 (10 µM) on ΔΨm was examined by the JC-1 assay in PM+H/R-treated cardiomyocytes. (N) The impact of Mdivi-1 on autolysosomes was examined by AO staining. (O) The effect of Mdivi-1 and Bafilomycin A1 (10 µM) on apoptosis by the TUNEL assay. (P, Q) The impact of Mdivi-1 and Bafilomycin A1 treatment on the expression of Drp1, Mff, LC3B, BNIP3, PUMA, p-p53, and BCL2 by Western blot. The data are expressed as the mean ± SEM (n=3-8). *P < 0.05 versus C, †P < 0.05 versus PM, ǂP < 0.05 versus H/R.

### PM+H/R significantly reduced miR-221/222 expression, while miR-221/222-enriched ADSC-Exo reduced PM+H/R-induced mitophagy and apoptosis

Our previous study showed that an abundance of miR-221/222 in ADSC-Exo reduced injury caused by I/R. ^22^ To explore whether miR-221/222 regulated PM+H/R-induced mitophagy and apoptosis, we assessed the levels of miR-221/222 using qPCR. Compared with the control group, the expression of miR-221/222 were significantly down-regulated in the PM or H/R treatment group. The levels of miR-221/222 in cardiomyocytes treated with PM+H/R were substantially lower than those in cardiomyocytes treated with PM or H/R alone (Figure 4A). However, the addition of ADSC-Exo increased the levels of miR-221/222 in PM+H/R-treated H9c2 cells (Figure 4B). Compared to H9c2 and ADSC cells, ADSC-Exo contained a large amount of miR-221/222 (Supplementary Figure 2). To elucidate the effects of miR-221/222 on mitophagy- and apoptosis-related proteins, we used “miRTarBase” target prediction and found that the 3’-UTR of LC3B, BNIP3, and PUMA contained the binding site program of miR-221/222 (Figure 4C). Our previous study demonstrated that miR-221/222 targets PUMA and regulates apoptosis. ^34^ To examine that miR-221/222 target 3’ UTR of LC3B and BNIP3, cells transfected with miR-221/222 mimics showed the reduced luciferase activity of the reporter genes (BNIP3 and LC3B). Additionally, no significant difference was observed in cells transfected with BNIP3/LC3B-MUT, suggesting that miR-221/222 can directly target BNIP3 and LC3B (Figure 4D). PM+H/R stimulation increased the mRNA expression and protein levels of BNIP3 and LC3B. However, ADSC-Exo or miR-221/222 mimics decreased the mRNA expression and protein levels of BNIP3 and LC3B (Figures 4E, 4F). However, compared with the PM+H/R+ADSC-Exo group, the expression of BNIP3 and LC3B increased in the PM+H/R+ADSC-Exo+miR-221/222 inhibitors group (Figure 4G). We also use the Seahorse analyzer to examine the efficiency of mitochondrial respiration and mitochondrial function. Treatment with the ADSC-Exo and miR-221/222 mimics increased OCR, ATP production, and maximal respiration in PM+H/R-treated cells, indicating that ADSC-Exo and overexpression of miR-221/222 provide protection against PM+H/R-induced mitochondria dysfunction (Figure 4H). MTT assay showed an increase in cell viability after treatment with ADSC-Exo and miR-221/222 mimics compared with PM+H/R, as shown in Figure 4I. We next investigated whether restoring miR-221/222 expression in the presence of PM+H/R could alleviate apoptosis. The TUNEL assay showed that ADSC-Exo administration and miR-221/222 mimics transfection significantly reduced PM+H/R- induced apoptosis (Figure 4J). However, miR-221/222 inhibitors increased the upregulation of apoptosis-related proteins (PUMA, p-p53, and cleaved caspase 3), whereas ADSC-Exo and miR-221/222 mimics protected against PM+H/R effect (Figures 4K, 4L). Additionally, to further test the roles of BNIP3 and LC3B in regulating apoptosis, we employed siRNA targeting BNIP3 and LC3B to knock down their expression. Through TUNEL assay and MTT assay, siBNIP3 and siLC3B reduced cell apoptosis and increased cell viability compared with the PM+H/R group (Figures 4M, 4I). Downregulation of BNIP3 and LC3B in PM+H/R-treated H9c2 cells reduced the expression of PUMA, p-p53, and cleaved caspase 3, while increasing the expression of BCL2 (Figures 4N, 4O). Notably, siBNIP3 and siLC3B effectively reduced the levels of apoptosis-related proteins but had no significant effect on fission-related proteins (such as Drp1 and Mff). Interestingly, siPUMA treatment had no significant impact on the levels of BNIP3, LC3B, Drp1, and Mff (Figure 4P). These results showed that ADSC-Exo administration reduced PM+H/R-induced mitophagy and apoptosis through the BNIP3/LC3B/PUMA pathway regulated by miR-221/222.

**Figure 4:**
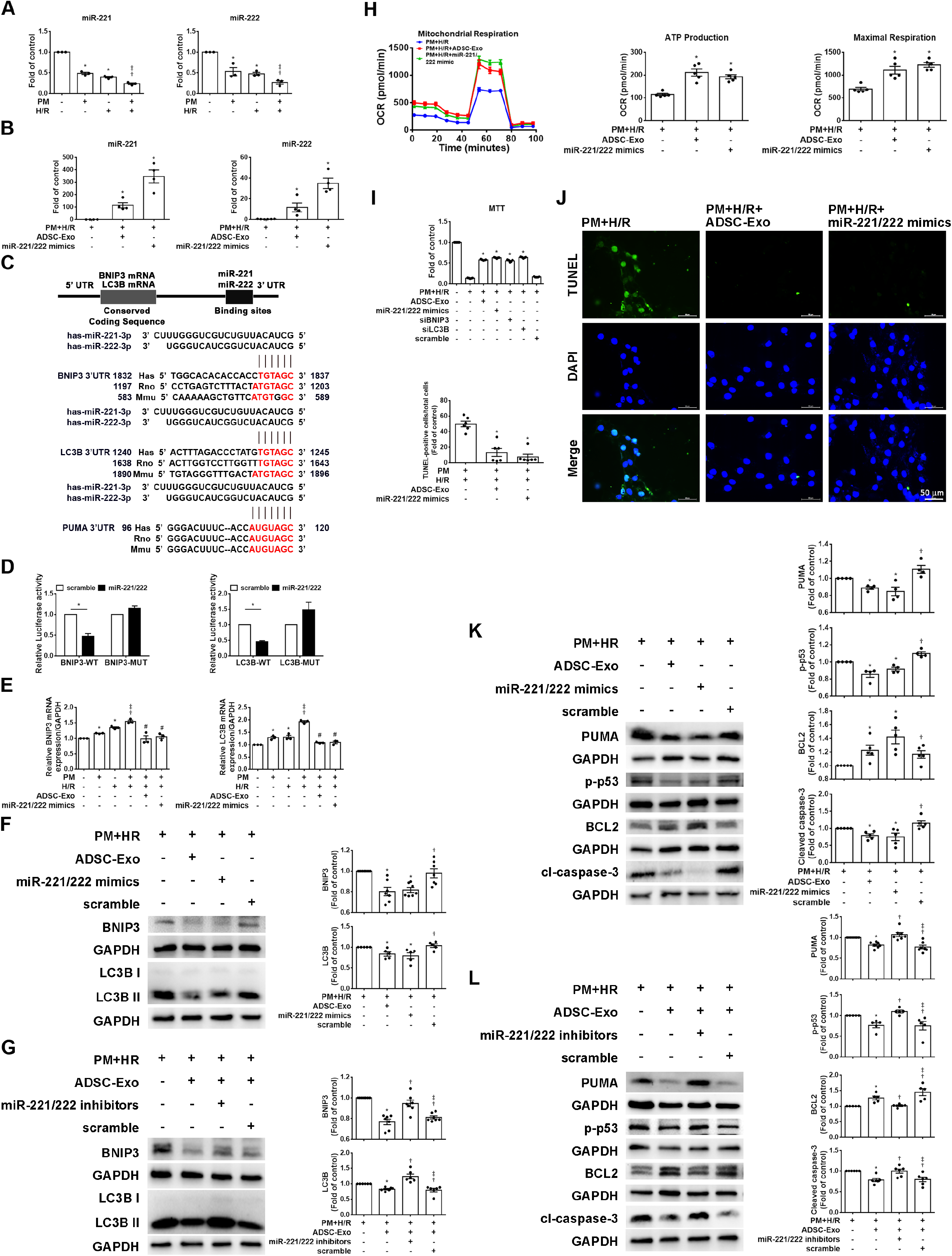
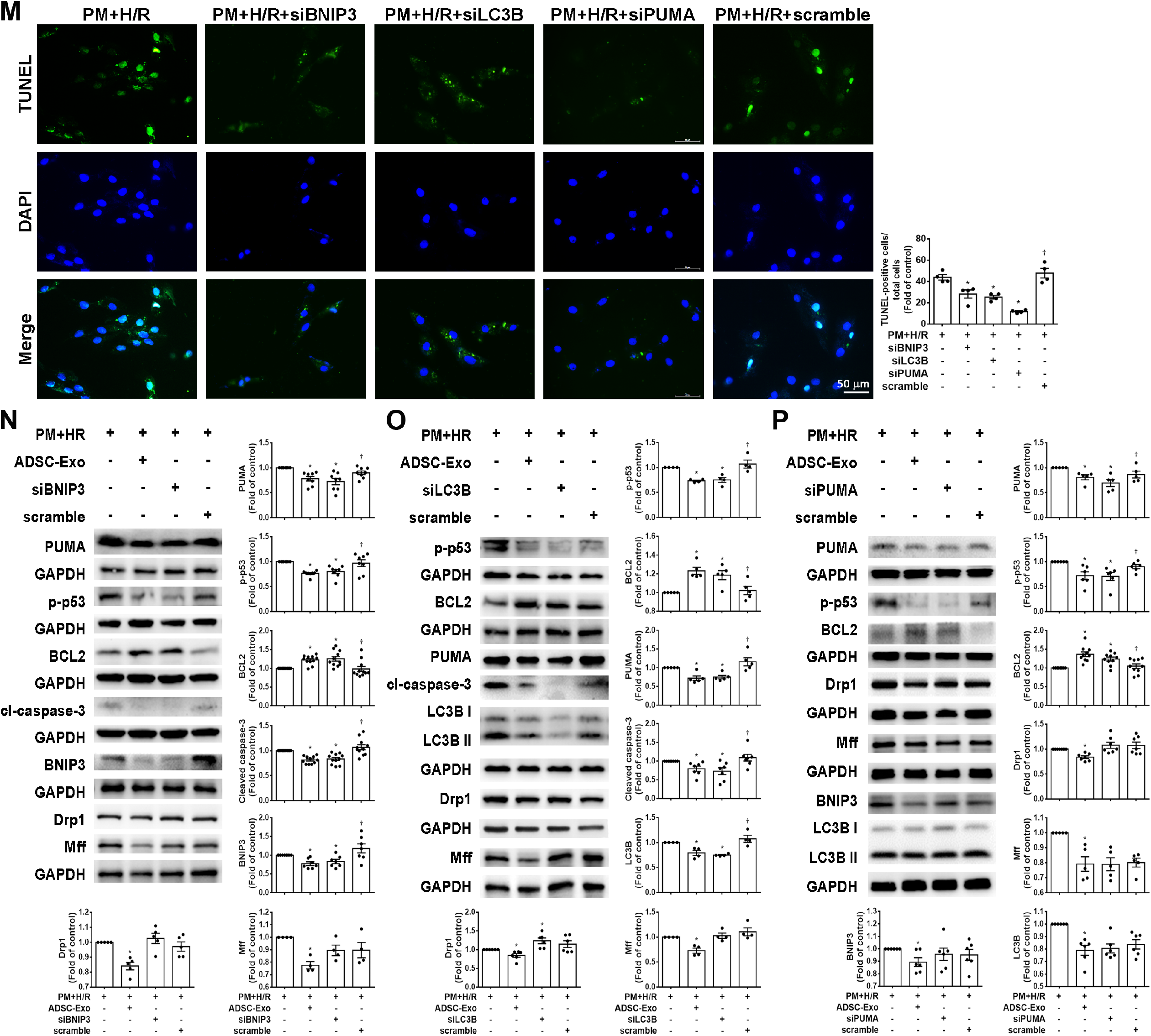
miR-221/222 regulates the expression of BNIP3 and LC3B, causing PM to aggravate myocardial H/R injury. H9c2 cells were transfected with miR- 221/222 mimics or inhibitors for 24 h, pretreated with PM for 6 h, and subjected to H/R for 6/12 h. (A) MiR-221/222 expression in various treatments was assessed using qPCR. (*P < 0.05 versus control, †P < 0.05 versus PM, ǂP < 0.05 versus H/R). (B) The impact of ADSC-Exo or miR-221/222 mimics on miR-221/222 expression in PM+H/R treatment was assessed by qPCR. (C) The schematic diagram depicted the binding of miR-221/222 to 3’ UTR of BNIP3, LC3B, and PUMA target genes. (D) Luciferase activities of BNIP3 and LC3B reporters were measured within the transfection of miR-221/222 mimics or scramble groups. (*P < 0.05 versus control). (E) Effects of ADSC-Exo or miR-221/222 mimics on BNIP3 and LC3B mRNA expression in PM+H/R-treated cardiomyocytes. (F, G) Effects of ADSC-Exo, miR- 221/222 mimics (F), and miR-221/222 inhibitors (G) on the expression of mitophagy-related proteins in PM+H/R-treated cardiomyocytes were detected by Western blot. (H) Effects of ADSC-Exo or miR-221/222 mimics on OCR in cardiomyocytes treated with PM+H/R. (I) ADSC-Exo, miR-221/222 mimics, siBNIP3, and siLC3B treatment on cell viability in PM+H/R-treated H9c2 cells as determined by the MTT assay. (J) The impact of ADSC-Exo and miR-221/222 mimics on apoptosis in PM+H/R-treated cardiomyocytes assessed via the TUNEL assay. Scale bar=50 μm. (K, L) The effects of ADSC-Exo, miR-221/222 mimics (K), and miR-221/222 inhibitors (L) on the expression of apoptosis-related proteins in cardiomyocytes exposed to PM+H/R were assessed by Western blot. (M) The effects of the downregulation of BNIP3, LC3B, and PUMA on cell apoptosis in PM+H/R-treated cardiomyocytes were examined by TUNEL assay. Scale bar=50 μm. (N, O) The effects of the downregulation of BNIP3 (N) and LC3B (O) on the expression of apoptosis-related proteins in cells exposed to PM+H/R by Western blot. (P) The effects of the PUMA downregulation on the expression of mitochondrial fission and mitophagy-related proteins in cells exposed to PM+H/R by Western blot. Data are expressed as mean ± SEM (n=4-11). *P < 0.05 versus H/R+PM, †P < 0.05 versus mimics or inhibitors or siRNA.

### ADSC-Exo attenuated PM+H/R-induced apoptosis by reducing ROS production, mitochondrial fission, and mitophagy

Mitochondrial ROS production was measured under ADSC-Exo, miR-221/222 mimics transfection, and MitoTEMPO conditions using MitoSOX Red staining. These treatments reduced the MitoSox Red fluorescence intensity in PM+H/R-treated cardiomyocytes assessed by fluorescence microscopy and flow cytometry (Figures 5A, 5B), indicating that ADSC-Exo and miR-221/222 mimics the reduction of mitochondrial ROS levels in PM+H/R-treated H9c2 cells. Next, JC-1 staining assessed by fluorescence microscopy and flow cytometry showed that PM+H/R treatment increased the number of low ΔΨm cells. Conversely, ADSC-Exo, miR-221/222 mimics, MitoTEMPO, or Mdivi-1 treatment reversed the collapse of ΔΨm, induced by PM+H/R (Figures 5C, 5D). H9c2 cells treated with ADSC-Exo, miR-221/222 mimics, MitoTEMPO and Mdivi-1 significantly increased ATP production in PM+H/R-treated cardiomyocytes (Figure 5E). The short mitochondrial length was observed in PM+H/R-treated cardiomyocytes, which was reduced by ADSC-Exo, miR-221/222 mimics, MitoTEMPO, and Mdivi-1 (Figure 5F). Western blot showed that the above treatment also significantly reduced the expression levels of Drp1 and Mff (Figure 5G). To further confirm the effects of ADSC-Exo and miR-221/222 mimics on mitophagy after PM+H/R injury, AO staining and RFP-GFP-LC3B staining were performed. PM+H/R treatment increased autophagosome presence, whereas treatment with ADSC-Exo, miR-221/222 mimics, MitoTEMPO, or Mdivi-1 reduced it (Figure 5H). RFP-GFP-LC3B showed autophagosomes and autolysosomes as yellow and red dots, respectively; ADSC-Exo and miR-221/222 mimics reduced mitophagy (Figure 5I). These results indicate that ADSC-Exo and miR-221/222 mimics reduce autophagosome presence. These treatments significantly reduced the levels of Beclin-1, BNIP3, and LC3B (Figure 5J). MitoTEMPO, Mdivi-1, and Bafilomycin A1 were used to examine the impact of ADSC-Exo and miR-221/222 mimics on apoptosis. These treatments significantly decreased apoptosis levels, measured using Annexin V-FITC/PI assays (Figure 5K). These treatments also reduced the apoptotic proteins PUMA and p-p53 while increasing BCL2 expression (Figure 5L). These results indicate that ROS triggers mitochondrial fission, mitophagy, and apoptosis in PM+H/R-treated cardiomyocytes, while ADSC-Exo inhibits ROS, mitochondrial fission, and mitophagy to attenuate cell apoptosis.

**Figure 5:**
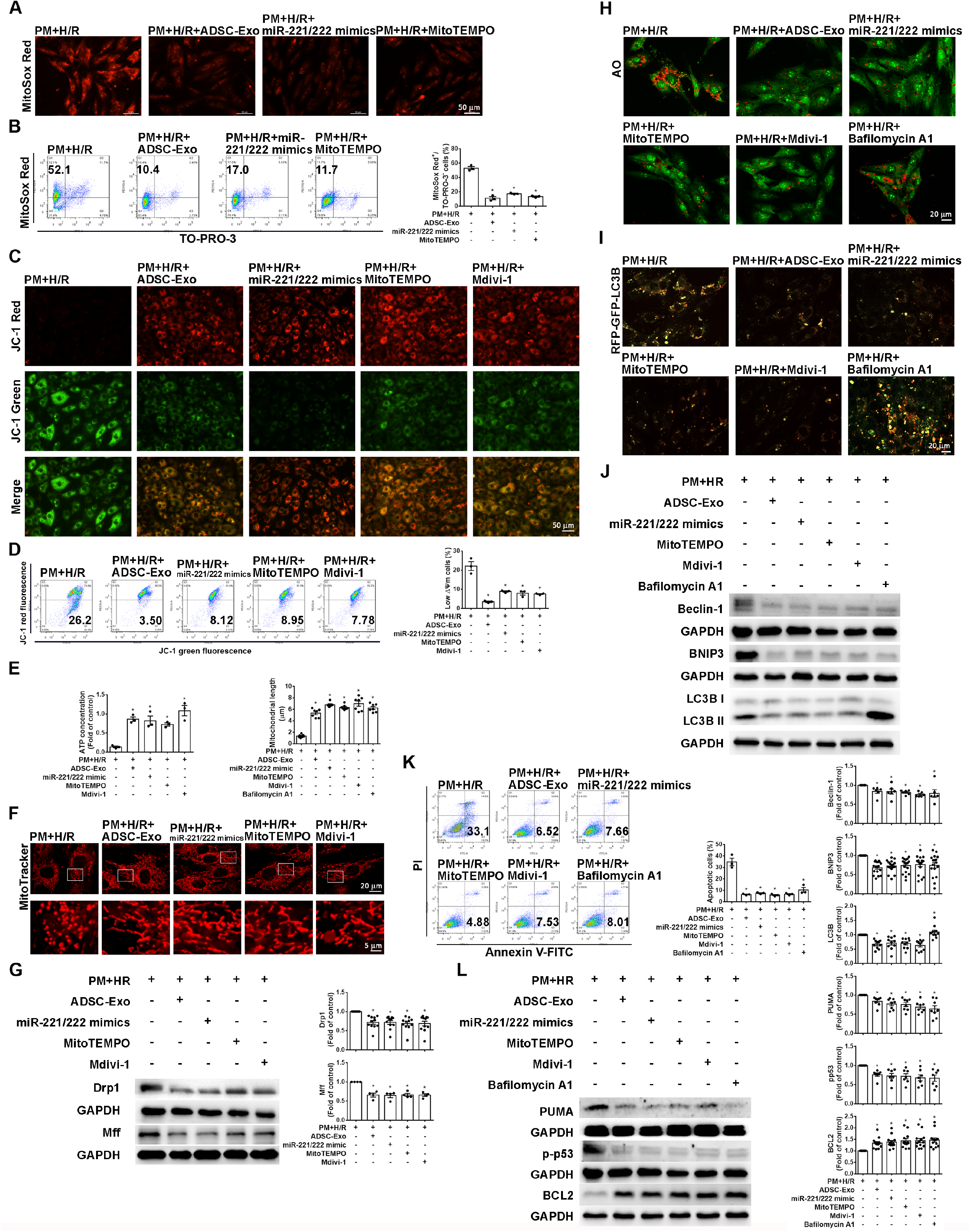
Abundant miR-221/222 in ADSC-Exo reduces mitochondrial ROS, mitochondrial fission, mitophagy, and apoptosis in cardiomyocytes exposed to PM+H/R. H9c2 cells were treated with ADSC-Exo (2 µg/mL) or transfected with miR-221/222 mimics for 24 h with PM (50 µg/mL) and subjected to H/R (6 h/12 h). (A, B) The effects of ADSC-Exo, miR-221/222 mimics, and MitoTEMPO treatment on mitochondrial ROS were evaluated by MitoSox Red staining under fluorescence microscopy and flow cytometry. The effects of ADSC-Exo, miR-221/222 mimics, MitoTEMPO, and Mdivi-1 treatment were assessed with (C, D) ΔΨm by JC-1 staining under fluorescence microscopy and flow cytometry, (E) ATP production using ATP assay, (F) mitochondrial length by the MitoTracker assay and (G) the expression of Drp1 and Mff by Western blot. (H) The effects of ADSC-Exo, miR-221/222 mimics, MitoTEMPO, Mdivi-1, and Bafilomycin A1 treatment on autolysosome accumulation using AO staining. (I) Cardiomyocytes were transfected with RFP-GFP-LC3 lentivirus for 72 h. Lentivirus allowed the distinction of autophagosomes (GFP+, RFP+, yellow plots) and autolysosomes (GFP−, RFP+, red plots) as the GFP fluorescence was quenched in the acidic autolysosomes. The effects of ADSC-Exo, miR-221/222 mimics, MitoTEMPO, Mdivi-1, and Bafilomycin A1 treatments on the formation of autophagosomes and autolysomes. (J) The effects of ADSC-Exo, miR-221/222 mimics, MitoTEMPO, Mdivi-1, and Bafilomycin A1 treatments on the expression of mitophagy-related protein levels were examined by Western blot. (K, L) ADSC-Exo, miR-221/222 mimics, MitoTEMPO, Mdivi-1, and Bafilomycin A1 effects on cell apoptosis were assessed by Annexin V/PI flow cytometry and Western blot. Scale bar=50 μm. Data are expressed as mean ± SEM (n=3-7). *P < 0.05 versus H/R+PM, †P < 0.05 versus mimics or inhibitors.

### MiR-221/222 attenuates PM+I/R-induced cardiac injury in WT and miR-221/222 KO mice and has the similar effect in transgenic (TG) mice

To elucidate whether ADSC-Exo protects mouse hearts from PM+I/R-induced damage through a miR-221/222-dependent mechanism. C57BL/6 (WT) and miR-221/222 KO mice were preconditioned with PM for 24 h. After 25 min of occlusion, followed by intramuscular injection of ADSC-Exo and miR-221/222 mimics into the anterior wall, and then 3 h of reperfusion. LDH and troponin I were significantly increased in the I/R and PM+I/R groups. ADSC-Exo or miR-221/222 mimics treatment significantly reduced PM+I/R-induced release of LDH and troponin I (Figure 6A). WT and miR-221/222 KO mice treated with ADSC-Exo or miR-221/222 mimics increased miR-221/222 expression levels under the PM+I/R environment (Figure 6B). ADSC-Exo or miR-221/222 mimics treatment increased EF and FS percentages in WT and miR-221/222 KO mice (Figure 6C). Therefore, we attempted to study the effect of ADSC-Exo on cardiac function by reducing ROS production. ADSC-Exo or miR-221/222 mimics significantly reduced cytoplasmic and mitochondrial ROS generation detected by DHE and MitoSox Red staining (Figures 6D, 6E). In addition, we observed that PM+I/R elevated the expression of BNIP3 and LC3B, while ADSC-Exo and miR-221/222 mimics reduced their expression by Western blot and immunohistochemistry (Figures 6F, 6G). Notably, a significant increase in PUMA, p-p53 expression, and a decrease in BCL2 expression after PM+I/R treatment were observed in WT and miR-221/222 KO mice, while ADSC-Exo or miR-221/222 mimics improved these changes (Figure 6H). Furthermore, ADSC-Exo or miR-221/222 mimics treatment reduced cardiac apoptosis, as demonstrated by the TUNEL assay (Figure 6I).

**Figure 6:**
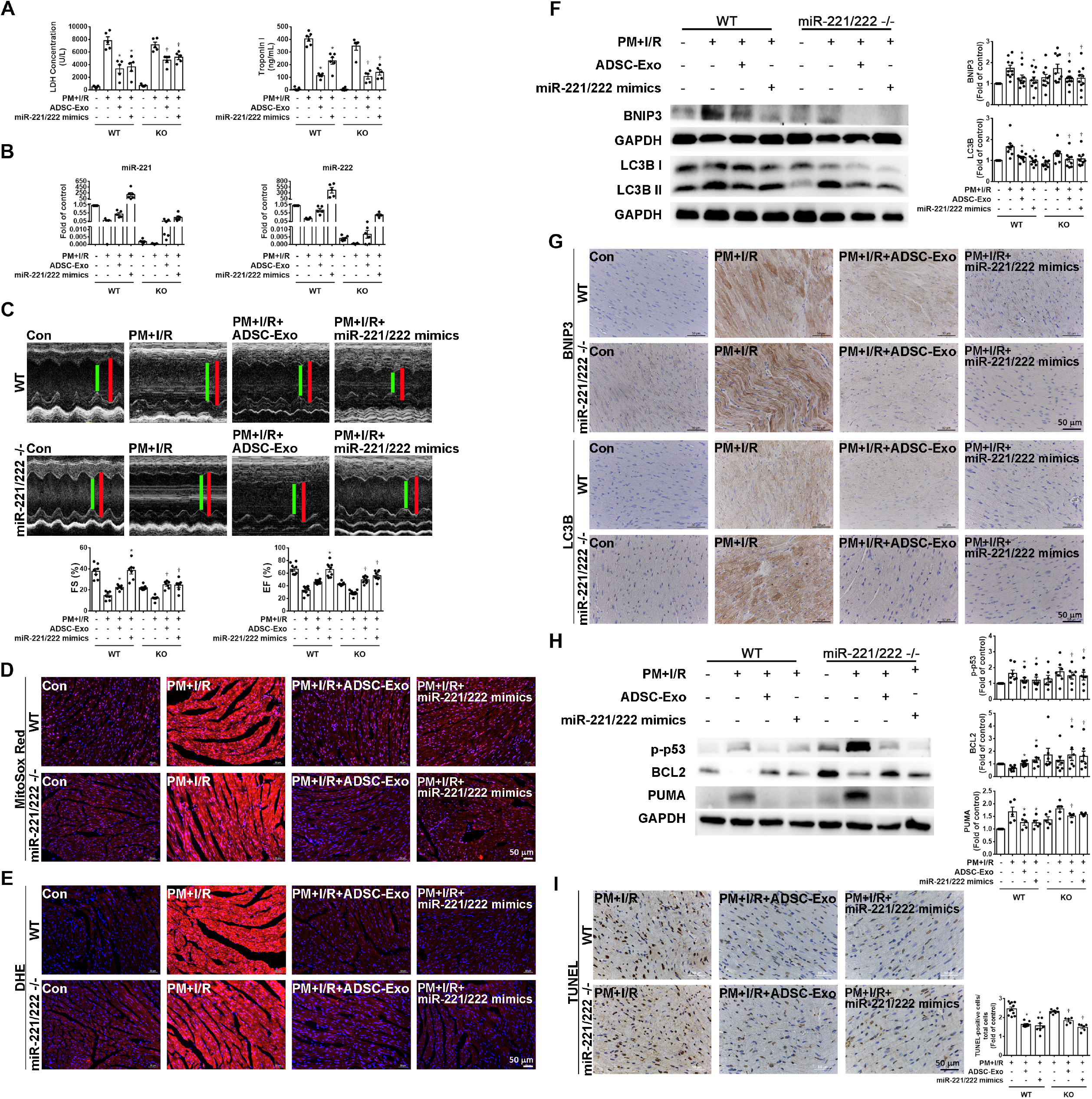
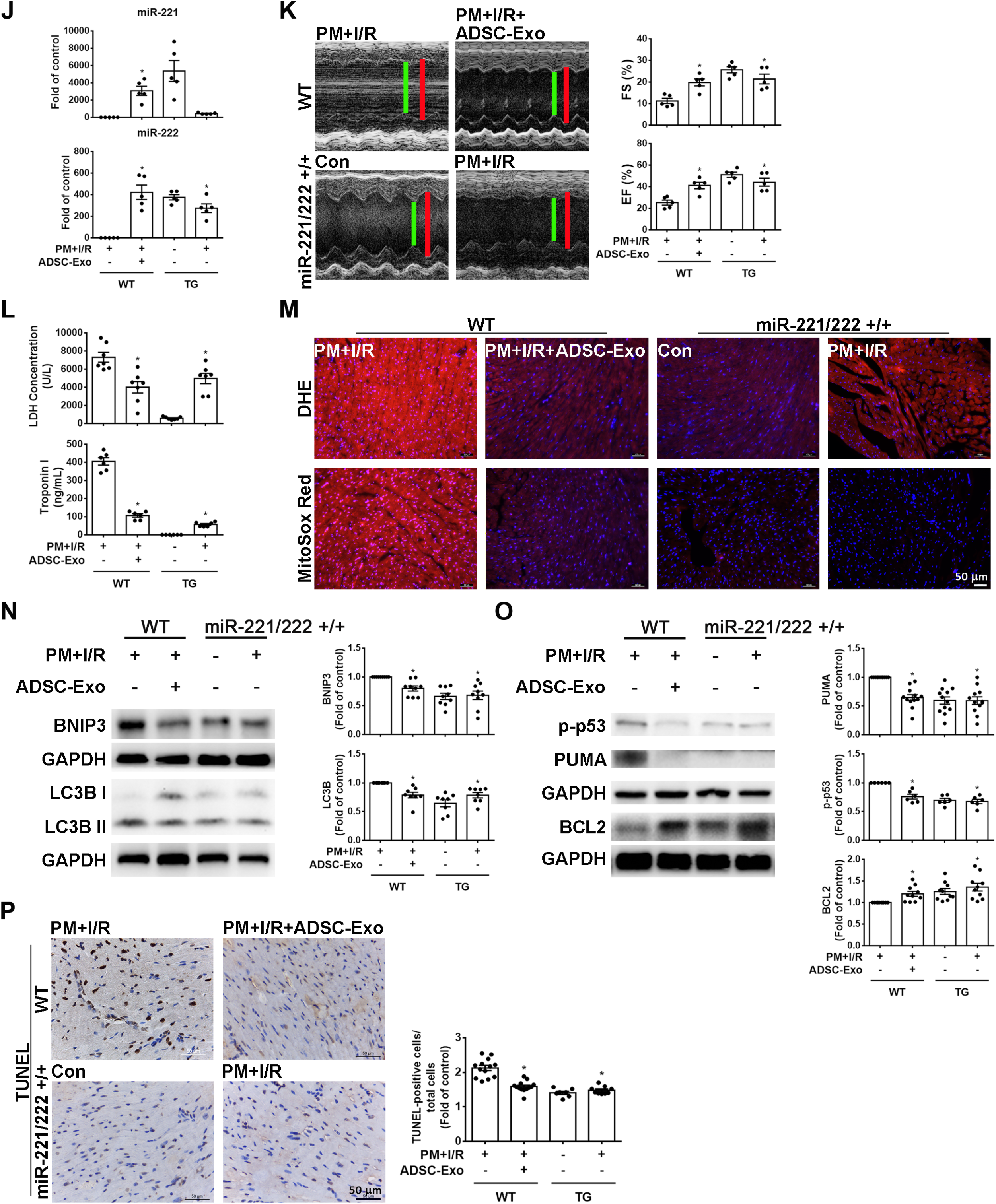
MiR-221/222 decreased PM+I/R-induced cardiac damage demonstrated by WT, miR-221/222 KO mice, and miR-221/222 TG mice. WT or miR-221/222 KO mice were preconditioned with PM for 24 h. Then, 25 min after occlusion, ADSC-Exo (100 μg protein in 50 μL) or miR-221/222 mimics (100 nM) were uniformly intramuscularly injected into the left ventricular marginal zone. The effects of ADSC-Exo and miR-221/222 mimics were assessed on various parameters: (A) levels of LDH and troponin I in plasma, (B) miR-221/222 expression by qPCR, (C) cardiac function by echocardiography, (D, E) ROS expression using MitoSOX Red staining and DHE staining, (F) levels of BNIP3 and LC3B expression by Western blot, (G) BNIP3 and LC3B expression assessed by immunohistochemistry, (H) expression of apoptosis-related proteins analyzed by Western blot, and (I) the number of apoptotic cells measured by TUNEL assay. To investigate the impact of miR-221/222 expression on PM+I/R-induced cardiac damage, miR-221/222 TG mice underwent preconditioning with PM for 24 h, followed by myocardial I/R, and were compared to PM+I/R WT mice and PM+I/R+ADSC-Exo WT mice. The following assessments were conducted: (J) qPCR to measure miR-221/222 expression, (K) echocardiography to assess heart function, (L) measurement of LDH and troponin I levels in plasma, (M) detection of ROS expression using MitoSOX Red staining. Additionally, the following assays were performed: (N) Western blot to examine mitochondrial autophagy-related protein levels, (O) Western blot to analyze apoptosis-related protein levels, and (P) TUNEL assay to quantify apoptotic cell numbers. Data are expressed as mean ± SEM (n=5-10). *P < 0.05 compared with WT PM+I/R, †P < 0.05 compared with KO or TG PM+I/R.

Importantly, we used overexpressing miR-221/222 TG mice as an animal model to further confirm the critical role of miR-221/222 in PM+I/R-induced cardiac injury. Meanwhile, myocardial miR-221/222 levels were significantly increased in miR-221/222 TG mice treated with PM+I/R compared with WT mice treated with the same treatment (Figure 6J). Echocardiography images showed that EF and FS were increased in PM+I/R-treated miR-221/222 TG mice compared with PM+I/R-treated normal mice, showing similar results to the ADSC-Exo-treated group (Figure 6K). Compared with the PM+I/R-treated WT group, the LDH and troponin I levels in PM+I/R-treated miR-221/222 TG mice were significantly reduced (Figure 6L). Furthermore, under PM+I/R treatment, miR-221/222 TG mice exhibited low ROS levels, and the results were similar to ADSC-Exo-treated PM+I/R-treated normal mice (Figure 6M). In addition, under PM+I/R treatment, miR-221/222 TG mice also showed low expression of BNIP3 and LC3B by Western blot, and the results were similar to ADSC-Exo-treated PM+I/R-treated WT mice (Figure 6N). Moreover, under PM+I/R treatment, miR-221/222 TG mice also showed low expression of PUMA and p-p53 detected by Western blot and reduced cell apoptosis detected by TUNEL. The results were consistent with ADSC-Exo-treated PM+I/R normal mice (Figures 6O, 6P). These findings strongly support that ADSC-Exo prevent myocardial PM+I/R injury through miR-221/222 *in vivo*. Taken together, these results indicate that I/R, combined with PM exposure, leads to increased ROS production, impaired mitochondrial function, maladaptive mitophagy, and ultimately increased apoptosis and cardiac damage.

## Discussion

Epidemiological studies have found a significant association between short-term exposure to environmental particulate matter (PM) and increased daily mortality and hospitalization from cardiovascular diseases. ^35,36^ In the present study, exposure to PM under I/R conditions exacerbated mitochondrial oxidative stress, leading to mitochondrial fission, mitophagy, and apoptosis. Our study shows that ADSC-Exo exerts its therapeutic effect by effectively alleviating the above effects, thereby mitigating mitochondrial dysfunction and cell apoptosis and ultimately restoring cardiac function. Importantly, our results provide evidence that miR-221/222 enriched ADSC-Exo reduces mitophagy and apoptosis through the BNIP3/LC3B/PUMA pathway and contributes to improving cardiac dysfunction.

Oxidative stress is essential to myocardial damage during acute I/R and chronic remodeling stages after myocardial infarction. ^37,38^ PM reduced cell viability and promoted apoptosis by increasing intracellular ROS production and activating MAPK and NF-κB signaling pathways in rat cardiac H9c2 cells. ^39,40^ Our current study shows that PM or I/R treatment administered individually significantly increases mitochondrial ROS and total ROS production, while PM exacerbates I/R-induced mitochondrial ROS and total ROS. Consistent with *in vivo* studies, PM exacerbates H/R-induced ROS production and apoptosis in cultured cardiomyocytes. Importantly, oxidative stress indicates mitochondrial abnormalities. ^37^ Mitochondrial quality control includes mitochondrial biogenesis, mitochondrial dynamics (fission/fusion), and mitophagy. ^41^ In rat aorta, PM increased SOD2 and mitochondrial fission proteins (Drp1 and Fis1) while reducing fusion proteins Mfn2 and OPA1 expression. ^18^ Previous studies have shown that mitophagy can clear damaged mitochondria after I/R injury, often viewed as a defensive or adaptive process. ^42,43^ However, excessive or uncontrolled mitophagy (maladaptive) may leave cells without enough healthy or functioning mitochondria to produce ATP, thus limiting cell viability. ^44^ Increasing evidence suggests that I/R may lead to an imbalance in the autophagy process, leading to additional cytotoxic damage that may lead to cell death. ^10,45^ On the other hand, PM triggered significant activation of mitophagy-related proteins in rat aorta, including LC3, p62, PINK, and Parkin. ^18^ Our current study demonstrates that PM+H/R decreases mitochondrial length and increases the expression of mitochondrial fission-related proteins Drp1 and Mff. Again, the combination of PM and H/R treatment increases the presence of autolysosomes and mitophagy-related proteins BNIP3 and LC3B expression. Furthermore, the degree of apoptosis is significantly reduced when mitochondrial ROS production, mitochondrial division, and mitophagy production are inhibited using MitoTEMPO, Mdivi-1, and Bafilomycin A1, respectively. These results support that mitochondrial oxidative stress is enhanced after PM+H/R treatment, leading to mitochondrial damage, increased mitochondrial fission, and improved mitophagy. A cascade of events driven by oxidative stress ultimately promotes apoptosis.

MiR-221/222 has been reported to be closely related to inflammation and the pathogenesis of cardiovascular diseases. ^25,46^ Our previous study showed that the expression of miR-221/222 in cardiac tissue was significantly reduced after I/R treatment. ^22^ Systemic inhibitions of miR-221/222 *in vivo* results in severe cardiac injury and prolonged viremia during viral myocarditis. ^47^ In this study, we found that combined treatment of I/R (H/R) and PM significantly reduced the expression of miR-221/222 compared with I/R (H/R) or PM alone. Our previous studies demonstrated that ADSC-derived conditioned media or exosomes containing large miR-221/222 can decrease I/R-induced cardiac injury. ^22,34^ Exosomes derived from various cells are considered a novel drug delivery method. For instance, Mesenchymal stem cells (MSC)-derived exosomes containing miRNA and proteins-are considered alternatives to cell therapy and play a promising role in cardiac regeneration and repair. ^48^ Few articles report the relationship between miR-221/222, mitophagy, and apoptosis. Elevated levels of miR-221/222 in glioblastoma promote cell survival while reducing these miRNAs leads to increased apoptosis through the upregulation of PUMA. ^49^ Another study in PTC cells showed that miR-221/222 reduced the expression of the autophagy protein LC3B and inhibited apoptosis. ^26^ MiR-221 significantly decreased cardiac H/R-induced autophagosome formation and reduced LC3 II/I ratio expression. ^50,51^ Extracellular vesicle-miRNA (has-miR221-3p) levels are downregulated after PM exposure and exhibit inflammation during pregnancy. ^52^ In this study, we demonstrated that the increase of miR-221/222 significantly reduced the expression of BNIP3, LC3B, and PUMA, indicating the regulatory role of miR-221/222 in mitophagy and apoptosis in PM+I/R (H/R)-induced cardiac injury.

Our study has thoroughly interpreted the regulatory role of miR-221/222 through the BNIP3/LC3B/PUMA pathway, but the two parts of the precise molecular mechanism require further study. First, verifying whether miR-221/222 regulates apoptosis through the AMPK/p27 pathway is necessary. This study showed that the AMPK/p27 pathway was induced under PM+I/R (H/R) conditions, whereas treatment with ADSC-Exo or miR-221/222 mimics reverted these changes. Furthermore, PM+I/R (H/R)-induced apoptosis was reversed when treated with pAMPK and p-p27 inhibitors (Supplementary Figure 3). Previous studies have shown that miR-221/222 inhibits p27 expression by targeting the 3’UTR. ^53,54^ ApoE inhibits smooth muscle cell proliferation by regulating p27 expression via miR-221/222. ^55^ Based on the above evidence, ADSC-Exo may reduce PM+I/R-induced apoptosis by targeting the miR221/222/AMPK/p27 signaling cascade. Secondly, how miR-221/222 regulates ROS production and mitochondrial fission remains to be determined. Previous studies have shown that miR-221/222 directly inhibits the expression of endothelial nitric oxide synthase (eNOS) or SOD2, reducing the production of nitric oxide (NO) and ROS, respectively. ^46,56^ Whether miR-221/222 regulates ROS production by targeting eNOS and SOD2 requires further study.

Overall, PM aggravates I/R (H/R)-induced cardiac injury. MiR-221/222-enriched ADSC-Exo reduces PM+I/R-induced mitophagy and apoptosis by downregulating BNIP3/LC3B/PUMA. This study provides a new treatment for PM+I/R-induced cardiac injury.

## Nonstandard Abbreviations and Acronyms

PM: particulate matter
I/R: ischemia/reperfusion
H/R: hypoxia/reoxygenation
ADSC-Exo: adipose-derived stem cells-exosome
miRNA: microRNA
BNIP3: Bcl-2 interacting protein 3
PUMA: p53 upregulated modulator of apoptosis
WT: wild type
KO: knockout
TG: transgenic
ROS: reactive oxygen species
3’UTR: 3’ untranslated region
MUT: mutation

## Article Information

## Acknowledgments

We thank the technical services provided by the “Transgenic Mouse Models Core Facility of the National Core Facility for Biopharmaceuticals, National Science and Technology Council, Taiwan” and the “Gene Knockout Mouse Core Laboratory of National Taiwan University Center of Genomic and Precision Medicine”. In addition, we thank the imaging core and flow cytometric analyzing and sorting core of the first core laboratory, National Taiwan University College of Medicine, for the technical support in image acquisition and analysis.

## Supplemental Material

Supplemental Expanded Materials and Methods

## Novelty and Significance section What is known?

- Epidemiology has previously demonstrated a strong relationship between exposure to particulate matter (PM) and cardiovascular disease.
- Myocardial I/R injury occurs when blood flow to the myocardium is temporarily blocked (ischemia) and then restored (reperfusion), which can lead to cardiac injury.
- MiR-221/222 present in ADSC-Exo that plays a role in the regulation of gene expression.

## What new information does this article contribute?

- Studies combining *in vivo* and *in vitro* models demonstrate that PM exposure aggravates cardiac injury by increasing ROS levels and causing mitochondrial dysfunction, ultimately leading to cardiomyocyte mitophagy and apoptosis.
- Transfection of wild-type mice and miR221/222 KO mice with ADSC-Exo and miR-221/222 mimics, as well as miR-221/222 overexpressing mice, significantly reduced PM+I/R-induced cardiac injury. At the same time, combined with in vitro experiments, it was detailed that this reduction was attributed to the ability of miR-221/222 to target and reduce the expression of BNIP3, LC3B, and PUMA, thereby regulating cardiomyocyte mitophagy and apoptosis.

This article provides novel insights into the relationship between PM exposure and cardiac injury during I/R and the potential therapeutic role of ADSC-Exo and miR-221/222 in alleviating these effects by modulating specific molecular pathways to modulate mitophagy and apoptosis.

